# Structural and biochemical analysis of a B12 super-binder

**DOI:** 10.1101/2024.02.16.580637

**Authors:** J.M. Martinez-Felices, J.J. Whittaker, D.J. Slotboom, A. Guskov

## Abstract

The gut microbiota is pivotal to human health, playing an important role in nutrient absorption and immune system. A prominent bacterial family in the human gut is *Bacteroidetes*. Cobalamin (Vitamin B12) is an essential micronutrient for these gut bacteria. Here, we focused on the BtuG family proteins in *B. thetaiotaomicron*, a model organism for this family, particularly the three homologs BtuG1, BtuG2, and BtuG3, which serve as cobalamin scavengers. Our study aimed to understand the structural and biochemical attributes of these proteins to describe their function in cobalamin acquisition. We solved the crystal structures of all three BtuG homologs bound to different cobalamin forms and the precursor cobinamide, and measured the binding kinetics by the grained coupled interferometry (GCI) technology. Our results reveal high binding affinities, in the low picomolar range underlining their critical role in cobalamin scavenging. Leveraging the high-resolution crystal structures, we successfully designed mutants of these BtuG homologs, resulting in both increased and decreased binding affinities for cobalamin and its precursor. This study furthers our understanding of bacterial adaptation mechanisms in the gut environment, and reveals a family of proteins with unique characteristics toward B12 scavenging, with implications for medical and biotechnological research.

## Introduction

The human gut is home to trillions of microorganisms, collectively known as the gut microbiota (Sender et al., 2016). The gut microbiota plays an essential role in host metabolism and immunity, and recent studies have highlighted the importance of the microbiota in maintaining human health (Clemente et al., 2012). The gut microbiota is a complex ecosystem composed of hundreds of species that interact with each other and with the host’s intestinal mucosal barrier (E. C. Martens et al., 2018; A. G. Wexler et al., 2018). The proportion of different micro-organisms inhabiting the human gut ranges widely between individuals. The *Bacteroidetes* phylum is one of the most abundant among the microbiota in human, accounting for more than 80% dominance in some individuals (Huttenhower et al., 2012).

Some essential vitamins are produced by the microbiota itself, such as vitamin K and certain B vitamins, which are then absorbed and utilized by the host (LeBlanc et al., 2013; Magnúsdóttir et al., 2015). These vitamins are also potentially shared among bacterial communities, thereby promoting microbial symbiosis (Ponomarova & Patil, 2015). On the other hand, vitamin B12 (cobalamin), must be obtained from animal-derived food sources (Martens et al., 2002) by both the human host and bacteria in the gut, leading to competition for the compounds (Degnan et al., 2014).

While humans absorb cobalamin from dietary sources in the small intestine, most of the cobalamin in the gut lumen is present in the form of inactive cobalamin analogues that cannot be used by the host nor the microbiota (Fang et al., 2017). Consequently, the gut microbiota has developed an intricate system to acquire cobalamin and transform it into biologically active forms that can be employed for growth and metabolic processes (Allen & Stabler, 2008; Degnan et al., 2014). Further evidence suggests that bacteria in the large intestine predominantly produce specialized cobalamin analogues, which may negatively impact host health and artificially increase measurements of circulating B12 in the blood (Allen & Stabler, 2008; Carmel, 2011).

To date, BtuB has been identified as the sole cobalamin transporter in the outer membrane (Heller et al., 1985), which enables the direct uptake of cobalamin from the extracellular environment (Cadieux et al., 2002). A broader range of membrane transport systems for cobalamin exist in the plasma membrane of both Gram-negative and Gram-positive bacteria. All characterized bacterial cobalamin transporters thus far belong to the adenosine tri-phosphate (ATP)-binding cassette (ABC) transporter superfamily (Nijland et al., 2022; Thomas & Tampé, 2020). Interestingly, soil bacteria were recently reported to encode a B12 transporter that additionally turns inactive cob(III)alamin analogues into cob(II)alamin coenzymes (Rempel et al., 2018). Additionally, the production of high-affinity cobalamin-binding proteins, known as corrinoid-binding proteins or cobalophores, further facilitates the sequestration and uptake of cobalamin (Degnan et al., 2014; A. G. Wexler et al., 2018).

In *B. thetaiotaomicron*, the cobalamin membrane transporter BtuB can be found in triplicates, where three BtuB homologs, BtuB1-2-3 have overlapping functions to provide efficient and competitive B12 transport (Degnan et al., 2014). The reasons for this seemingly inefficient use of genetic space are not completely clear, but *Bacteroides* species have been described as genetic “pack rats”, bearing multiple paralogous groups of genes, and therefore will not need to rely on unpredictable mutations (Wexler, 2007). Knockout competition studies showed that BtuB2 provides the highest fitness of all three BtuB homologs. However, BtuB1 and BtuB3 can provide specific fitness to different variants of cobalamin in absence of BtuB2 (Degnan et al., 2014).

The BtuG family of cobalamin scavengers, present exclusively in *Bacteroidetes*, is situated in genetic operons controlled by B12-activated riboswitches. Nearly all the members of *Bacteroidetes* gut bacteria encode genes for BtuG homologs (Wexler et al., 2018). The three genetically distinct BtuG homologs in *B. thetaiotaomicron* are each encoded in a genetic *locus* along one of the three genes encoding BtuB homologs (Wexler et al., 2018). BtuG is a cell-surface exposed lipoprotein that binds cobalamin with high affinity and associates with BtuB to support efficient uptake of B12 (Wexler et al., 2018). Although BtuB is capable of importing cobalamin on its own, BtuG-mediated cobalamin scavenging is critical for *in vivo* growth of *B. thetaiotaomicron* in germ-free mice (Degnan et al., 2014). Notably, BtuG can extract cobalamin from the human carrier protein intrinsic factor directly from the organism’s surface or in the membrane of bacterial extracellular vesicles that contain BtuG proteins, providing a competitive advantage for cobalamin sources in the human gut (Juodeikis et al., 2022; A. G. Wexler et al., 2018).

In this study, we report the crystal structures of the three BtuG homologs present in *B. thetaiotaomicron*: BtuG1, BtuG2 and BtuG3; bound to two cobalamin variants: cyanocobalamin (CNCbl) and hydroxycobalamin (OHCbl); and the cobalamin precursor Cbi. We measured binding kinetics for all three homologs for their substrates using GCI technology, revealing that all of them bear extremely high affinity for cobalamin variants, in the low picomolar range, and medium affinities for the precursor Cbi in the mid-micromolar range. We found up to 100-fold differences in affinity among different homologs for Cbi. Moreover, supported by structural and biochemical data, we designed single-mutants that achieved both increase and decrease of affinities for their substrates. Overall, these results reveal the molecular mechanisms underlying cobalamin scavenging by *B. thetaiotaomicron*.

## Results

### Structures of BtuG homologs BtuG1, 2 and 3 bound to different cobalamin variants

BtuG2 from *B. thetaiotaomicron* has been proposed to be a cobalamin scavenger (Wexler et al., 2018). We aimed to further study the role of BtuG2 as a cobalamin scavenger *in vitro* and compare its properties with two homologs from the same organism, BtuG1 and BtuG3. In order to produce the three homologs as soluble proteins in *E. coli*, the native lipoprotein export sequences were removed from each BtuG homolog (Fig. 2a), leading to heterologous expression in the cytoplasm instead of targeting the outer leaflet of the outer membrane. For clarity, residues are numbered based on the native protein sequence, including the lipoprotein export sequences.

Based on sequence alignment, BtuG2 and BtuG3 share the highest similarity with ∼88% (∼79% identity), (Fig. 2d), while BtuG1 shares lower similarity with BtuG2, ∼40% (∼24% identity) (Fig. 2c). While BtuG2 and BtuG3 sequences contain only the predicted β-propeller domain, BtuG1 has an additional C-terminal domain that has been identified as a homolog of BtuH (Putnam et al., 2022), a cobalamin scavenger present as an independent gene in the cobalamin transporter *loci* 2 and 3 of *B. thetaiotaomicron*, but fused to BtuG1 in *locus* 1 (Putnam et al., 2022). Although BtuH has been reported as a cobalamin scavenger and it was expressed along BtuG1, the predominant purified protein presents a molecular mass of ∼50 kDa in SDS-PAGE gel (Suppl. Fig. 1), lower than the expected mass of ∼70 kDa of the BtuG1 and BtuH domains together. We hypothesize the overexpressed protein undergoes proteolytic cleavage of ∼20 kDa in the c-terminal corresponding to the BtuH domain, given that the construct contains an N-terminal histidine tag used for affinity purification. Mass spectrometry analysis of the BtuG1 SDS-PAGE band determined a main population of 46,795 Da, corresponding to a ∼23 kDa cleavage from the construct sequence (Suppl. Fig. 2). This cleavage could occur during either protein expression or purification. Sensitivity towards cleavage has been already reported for BtuH2 (Putnam et al., 2022). Moreover, no BtuH densities could be identified in either of the two BtuG1 crystal structures that we determined. Most of BtuH binding site contacts to cobalamin involve residues from the last 20 kDa of the protein. Therefore, we consider our kinetic results to be the contribution of solely BtuG1.

**Figure 1.**
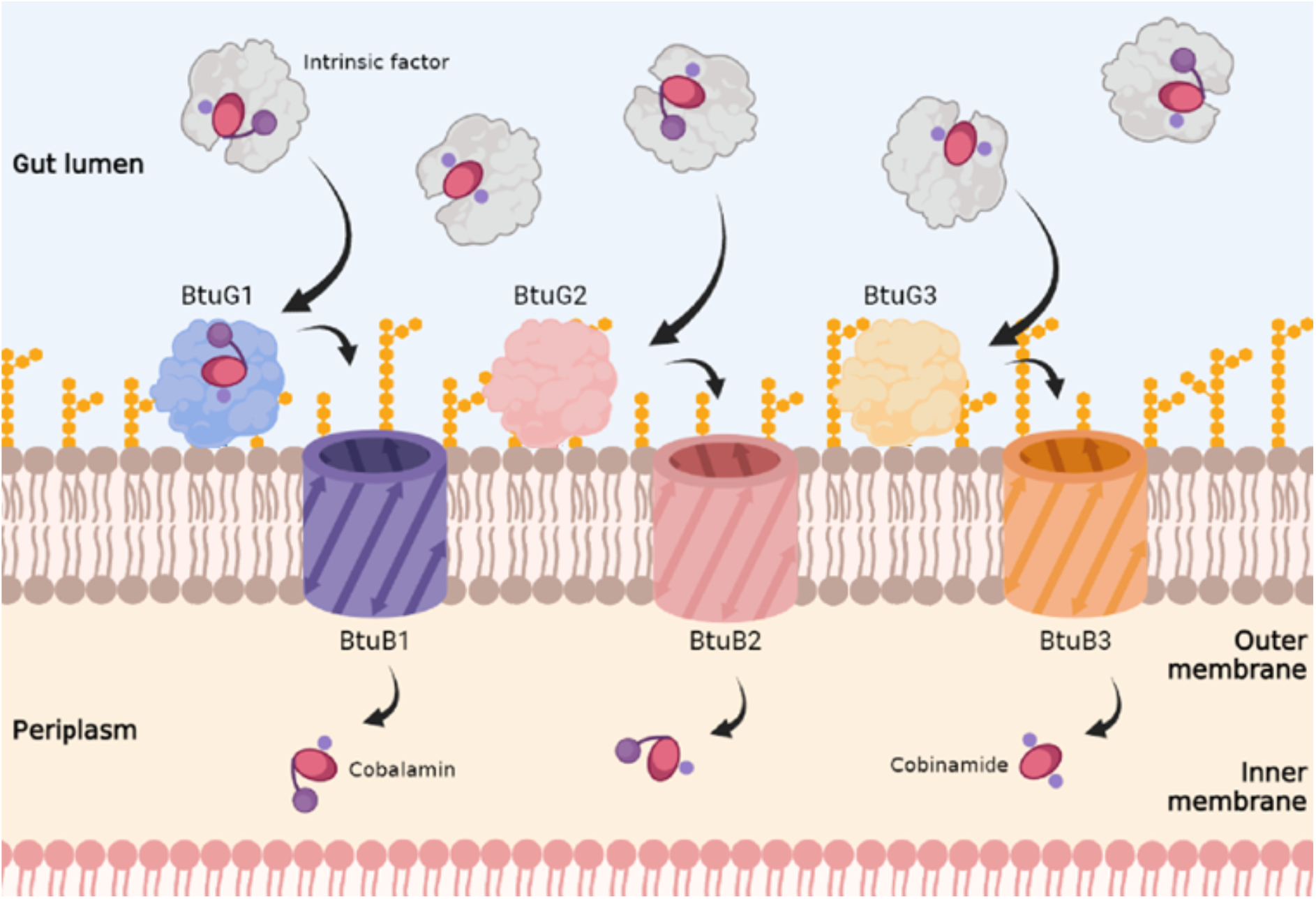
Scheme of redundant vitamin B12 uptake by *Bacteroidetes thetaiotaomicron.* *B. thetaiotaomicron* encodes three operons encoding homologous BtuG-BtuB pairs, providing three similar transport routes for cobalamin, its variants and structurally related precursors (Cbi). Vitamin B12 in the form of cobalamin and Cbi molecules (red ovals for the corrin ring, small blue ball for the functional group coordinating the cobalt ion and purple ball for the DMB group) are transported through the upper gut attached to a protective intrinsic factor molecule of human synthesis (grey globule).

**Figure 2.**
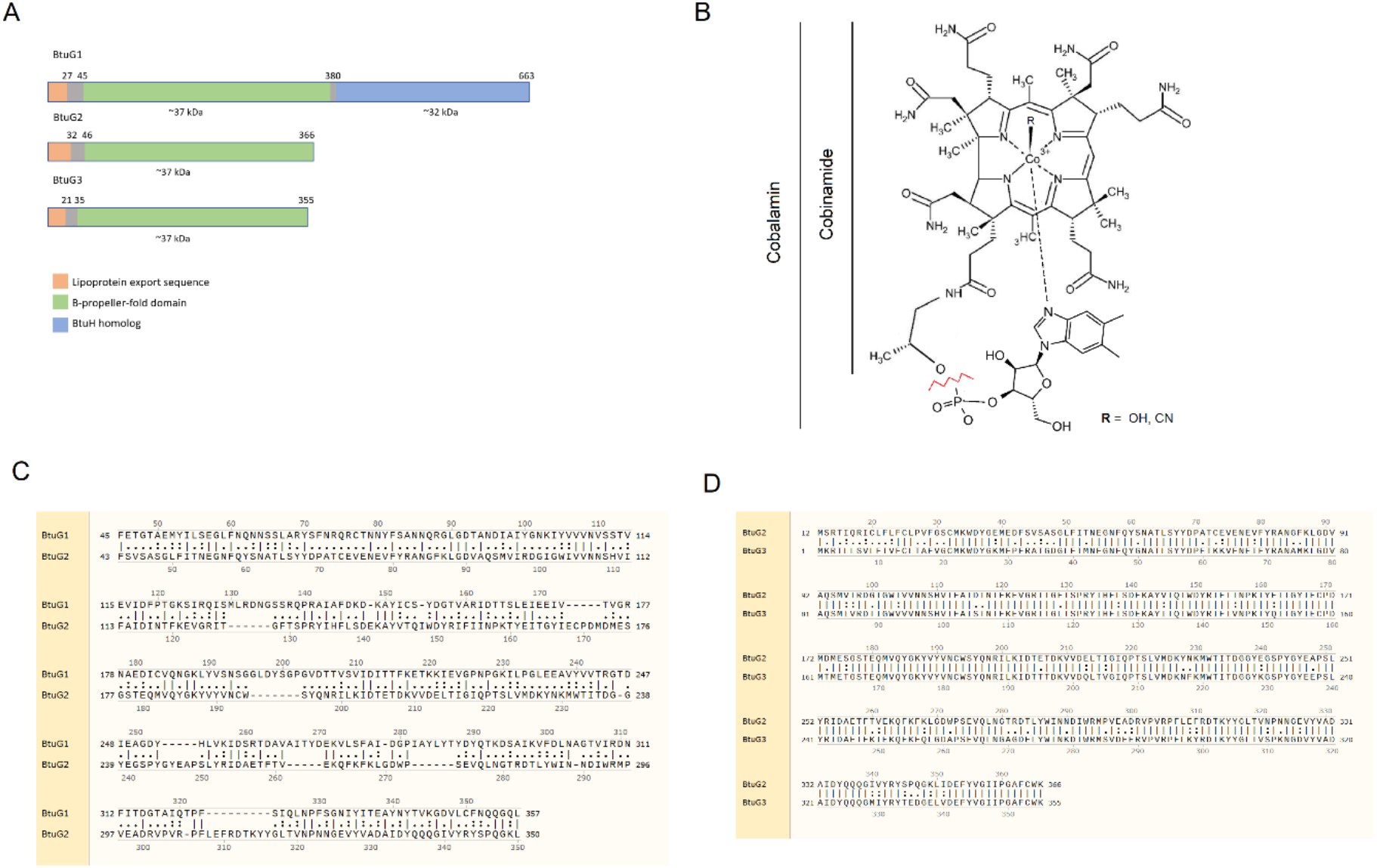
(A) Sequence homology of BtuG1-2-3. (B) Structural diagram of cobalamin and Cbi. The red line indicates where Cbi is truncated compared to Cbl. Sequence alignments of BtuG2 against (C) BtuG1 and (D) BtuG3. Lines between both sequences represent identical residues, two dots for similar and one, not similar.

We now report the structure of BtuG2 in complex with two cobalamin variants, CNCbl and OHCbl, and the cobalamin precursor Cbi, at 2.0 Å, 1.50 Å and 1.69 Å, respectively; revealing a 7-bladed β-propeller fold with two openings at each side of the protein. The BtuG2 conformations are identical regardless of which substrate was bound, with a RMSD of 0.074 Å between BtuG2-CNCbl and BtuG2-OHCbl; and 0.125 Å RMSD between BtuG2-CNCbl and BtuG2-Cbi. For clarity, we will base our analysis on BtuG2-CNCbl structure, since it is the structure with the highest resolution that has all protein-substrate interactions. All variants of B12 solved in complex with BtuG2 bind in the same aperture of the β-propeller-fold of BtuG2, which bears a high density of negative charges (Wexler et al., 2018).

At the binding site, multiple BtuG2 residues make contacts with the substrate: ten hydrophobic amino acid residues stabilize the hydrophobic moieties of the substrate (Fig. 3b, Suppl. Fig. 3b), while eight polar side chains, with the main chain of a glycine residue form electrostatic contacts with the vitamin (Fig. 3a). Hydrophobic interactions in BtuG2 are dominated by aromatic sidechains of tryptophan, tyrosine and phenylalanine, that form an envelope around the corrin ring of Cbl and Cbi and are located facing the hydrophobic regions of the ring. Two isoleucines, Ile148 and Ile359, are the only non-aromatic hydrophobic interactions, facing the hydrophobic chain of two amide sidechains (Suppl. Fig. 3b). In addition, Phe58 makes an edge-to-face aromatic-aromatic contact with the DMB moiety of CNCbl and OHCbl (Fig. 3b, Phe58), where aromatic rings are rotated 90°, similar to the interaction between Phe401 and CNCbl in BtuH2 (PDB: 7BIZ, Putnam et al., 2022). Polar interactions target the corrin ring amide sidechains, which bear both positive and negative dipoles, allowing for a variety of interactions and providing specificity. All six sidechains are involved in at least one interaction with a BtuG2 residue. Asn107, Gln93, Gln219, Ser131, Ser195, Trp149 and Gly56 make hydrogen bonds with the substrate to stabilize the corrin ring. Moreover, Glu55 and Arg135 sidechains, bearing net negative and positive charges, respectively, are involved in strong electrostatic interactions with the ligand (Sippel & Quiocho, 2015).

**Figure 3.**
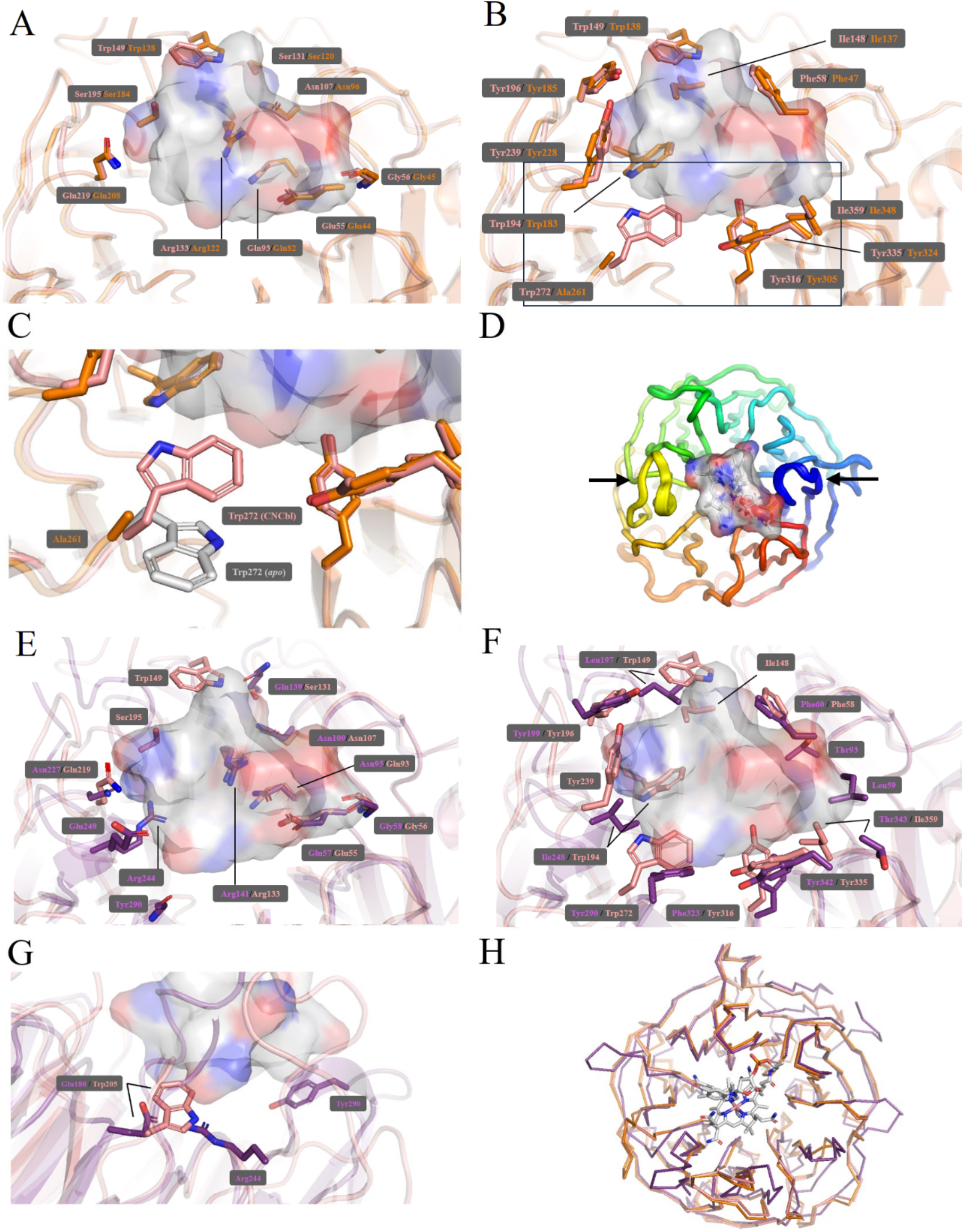
Structural insights into BtuG homologs from *B. thetaiotaomicron*, bound to Cbl and Cbi. Comparison between the binding site polar contacts (A and E) and hydrophobic interactions (B and F) of BtuG2 (pink), with BtuG3 (orange) and BtuG1 (Purple) toward CNCbl (grey surface, from BtuG2 structure). Protein backbone is represented in cartoon with interacting residues in sticks. Blue colour for nitrogen and red for oxygen. Black frame in (B) indicates the zoomed region for (C), comparing the position of Trp272 in *holo-*BtuG2 (pink) and *apo-*BtuG2 (grey sticks). (D) Rainbow-colored morph structure of the conformational changes between *apo*-BtuG2 and *holo-*BtuG2. (G) Same color-coded as in (E) and (F), purple sticks represent BtuG1 sidechains of residues mutated to tryptophan and in pink sticks, the reference tryptophan from BtuG2. (H) Overall conformation of the three BtuG homologs, with same color-coding as in (A) and (E), with CNCbl from BtuG2 as a reference. Protein backbone is represented in ribbon.

The crystal structure of the soluble domain of BtuG2 from *B. thetaiotaomicron* was previously solved in the *apo*-state, (PDB ID: 3DSM). In BtuG2-CNCbl, we observe subtle embracing of the cobalamin molecule by loop 1, connecting β-sheets 2 and 3 from blade 1; and loop 17, connecting β-sheets 2 and 3 from the opposing blade 5 (Fig. 3d). However, the overall conformation remains intact, with a RMSD of 0.29 Å.

To study the role of the homologs BtuG1 and BtuG3, we solved the structure of BtuG1 in complex with CNCbl and Cbi (PDB ID: 8PTD and 8PTF, respectively; Fig. 3e, f and h), and the structure of BtuG3 in complex with Cbi (PDB ID: 8PUJ, Fig. 3a, b, c and d). Their structures are practically identical with a RMSD of 0.42 Å (Fig. 3H). The BtuG3 binding site provides the same sidechain contacts as BtuG2 except for Trp272, which in BtuG3 is substituted by an Ala261 (Fig. 3a, b and c, Suppl. Info. 3b), thereby removing the indole hydrophobic stabilization of the corrin ring of Cbi.

Despite the relatively low sequence similarity, BtuG1 and BtuG3 possess identical folds, with a RMSD of 1.5 Å (Fig. 3H) and share multiple binding contacts to CNCbl and Cbi with BtuG2 (Fig. 3e and f, Suppl. Fig. 3). The binding site of BtuG1 provides ten hydrophobic interactions to stabilize the hydrophobic moieties of Cbl. Similar to BtuG2, these residues are distributed around the corrin ring (Fig. 3e). In the BtuG1 binding site, four aromatic residues and five non-aromatic five, interact with the substrate (Suppl. Fig. 3a). An edge-to-face aromatic interaction by Phe60 stabilizes the DMB moiety of cobalamin, just as in the case of BtuG2 and 3 (Fig. 3b). Among the ten residues that make polar contacts within the BtuG1-Cbl interface, there is a higher number of ion-dipole interactions, provided by Arg141, Arg244, Glu139 and Glu249. Six other residues make hydrogen bonds. Notably, Tyr290 is involved in a hydrophobic interaction at its aromatic ring, while its backbone oxygen is the receptor of a hydrogen bond. Overall, ∼60% of these contacts are similar or identical to BtuG2.

### All BtuG homologs bear extreme affinities toward cobalamin variants

We measured the affinities of BtuG1, 2 and 3 toward CNCbl, OHCbl and Cbi using GCI technology (Table 2, Suppl. Fig. 4, Patko et al., 2012). Each homolog was attached to a chip with Nickel-NTA surface via an engineered histidine tag after purification and the binding kinetics toward CNCbl, OHCbl and Cbi were measured at different substrate concentrations (Table 2, Suppl. Fig. 4). All three BtuG homologs showed higher affinities toward cobalamin variants CNCbl and OHCbl than for the precursor Cbi. BtuG1, 2 and 3 have affinities in the 1-10 picomolar range for both CNCbl and OHCbl (Table 2).

Surprisingly, the affinities for Cbi exhibit the highest variation between the three homologs. The Cbi dissociation kinetics can only be described when two populations (with faster and slower *k*_off_), are modelled together (heterologous ligand), leading to two dissociation constants: 5.8·10^-5^ M and 1.4·10^-5^ M; 4.3·10^-7^ M and 5.9·10^-8^ M; and 3.1·10^-5^ M and 3.9·10^-6^ M for BtuG1, 2 and 3, respectively, with the affinity for BtuG2 ∼100-fold higher than BtuG3 and BtuG1 (Table 2). The presence of two kinetic affinities for BtuG proteins has not been described before, and seems to be exclusive to Cbi, as it is not observed for CNCbl and OHCbl. Whether this is just a limitation of our experimental setup or might have some functional implication remains to be explored.

### BtuG2 mutants have its affinity toward Cbi decreased

In order to explore the role of individual residues involved in the BtuG2:CNCbl complex formation we mutated residues Glu55, Phe58, Gln93, Gln219 and Trp272 (Fig. 3a and b); substituting them for an alanine residue. The rationale for mutagenesis is that Glu55, Gln93 and Gln219 make mono-or bidentate electrostatic contacts with the substrate (Fig. 3a) hence their removal is expected to weaken BtuG2 affinity toward Cbl and Cbi. Phe58 is involved in an aromatic interaction with DMB moiety, hence relevant for Cbl variants but not Cbi (Fig. 3b). Lastly, Trp272 was found to be the only residue to change its conformation upon substrate binding, flipping its sidechain to face the corrin ring of the substrate (Fig. 3c).

Affinities of all BtuG2 mutants for CNCbl and Cbi were tested using GCI (Table 2). While dissociation constants for CNCbl remained in the low picomolar range, the affinity of BtuG2 mutant W272A for Cbi was weaker by more than 10-fold (Table 2), likely due to the loss of indole sidechain group of tryptophan which has been reported to stabilize the corrin ring of cobinamide (Mireku et al., 2017).

### BtuG1 and BtuG3 mutants can modulate their affinity toward cobalamin

BtuG2 has 100-times higher affinity for Cbi than BtuG1 and BtuG3 (Table 2). In an effort to modulate the affinities of BtuG1 and BtuG3 for the substrates, we made one BtuG3 and three BtuG1 mutants with the aim to increase the stability of the corrin ring of both Cbl and Cbi. Ala261 is the only residue that is different in the BtuG3 binding site compared to the binding site of BtuG2 (Fig. 3c). Therefore, we designed the A261W mutant, mimicking the native binding site of BtuG2. The affinity for Cbi was increased 10-fold, from 3.1·10^-5^ M in the WT to 4.3·10^-6^ M in the A261W mutant, at least for one of the binding components revealed by the heterogeneous kinetics. The affinity for Cbl remained the same.

In BtuG1, E180, R244 and Y290 were mutated to a tryptophan that provides the hydrophobic interaction present in BtuG2 via Trp194 but is missing in BtuG1 (Fig. 3e). No significant affinity increase towards Cbi was detected. Instead, E180W and R244W had significant decreases in the affinity for CNCbl and OHCbl (Table 2), most likely caused by the perturbations in the binding site, potentially impeding the correct interaction with the substrate (Fig. 3G).

## Discussion

In this study, we have determined the crystal structures of the three BtuG homologs from *B. thetaiotaomicron*, BtuG1, BtuG2 and BtuG3; bound to three of their substrates, namely CNCbl, OHCbl and Cbi at high resolution, between 1.5 and 2.5 Å (Fig. 3, PDB IDs: BtuG1-CNCbl: 8PTD, BtuG1-Cbi: 8PTF, BtuG2-CNCbl: 8BAI, BtuG2-OHCbl: 8BB0, BtuG2-Cbi: 8BAZ, BtuG3-Cbi: 8PUJ). In total, these six crystal structures allowed us to perform the detailed analysis of the interactions between the BtuG family of proteins and vitamin B12. Binding kinetics for these interactions were measured using GCI technology. Previously, the crystal structure of BtuG2 in the *apo-*form showed a protein topology of a 7-bladed β-propeller fold (PDB ID: 3DSM). Remarkably, BtuG2 does not undergo major conformational changes upon substrate binding (Fig. 3d), with only loop 1 and loop 17 slightly approaching the Cbl substrate, with an overall RMSD of 0.29 Å. We, therefore, conclude that the low-picomolar affinity of all BtuG homologs for Cbl (Table 2) is the result of a highly tailored binding site, involving up to 37 protein:ligand contacts, and not from the large conformational changes. Extremely high affinities (low picomolar to femtomolar) usually involve protein conformational changes that ‘trap’ the substrate, as seen in the case of streptavidin binding biotin (Weber et al., 1989). On the other hand, other proteins present in nature with highly tailored binding sites that do not undergo conformational changes fall short on affinity compared to BtuG, e.g., antibodies and nanobodies (Huo et al., 2020; Muyldermans & Lauwereys, 1999); and the neurexin-neuroligin complex (Leone et al., 2010; Südhof, 2008). Therefore, the BtuG:Cbl complex is of extraordinary strength, placing BtuG within the “Superbinder” affinity class, previously reserved only to engineered proteins with femtomolar affinities (Jonsson et al., 2008; Zhou et al., 2014) and it makes its ‘static’ binding mode unique among known substrate scavengers. The nature of the ligand and the vast number of interactions are the main drivers of such an extreme affinity. Recently, the crystal structures of BtuG2 and BtuG3 bound to different Cbl variants, and the Cryo-EM structure of BtuG1 in complex with BtuB1 were reported (Abellon-Ruiz et al., 2023). The reported structures and their analysis of the binding interactions are consistent with the structures provided in this study, reinforcing the conclusions made regarding the interaction with the substrate and the binding mode to cobalamin variants.

The redundant outer membrane cobalamin transport in *B. thetaiotaomicron* consists of three homology groups of one scavenger (BtuG) and one outer membrane transporter (BtuB, Fig. 1). The contribution of each BtuB homolog to the organism fitness was found crucial for transport of different cobalamin variants (Degnan et al., 2014). However, in that study the contribution of BtuG as a scavenger was not addressed. BtuG1-2-3 share the same topology (Fig. 3H). BtuG2 is most similar to BtuG3 with an 84% similarity and 0.42 Å RMSD. In contrast, BtuG1 has a higher degree of variability at the sequence level compared to its BtuG homologs, with a similarity of 24%. Therefore, there are multiple interactions that are not conserved, compared to BtuG1-2 (Fig. 3e, f and g). Our kinetic studies reveal that the three BtuG homologs bind the cobalamin precursor Cbi with different affinities. The highest affinity was found for BtuG2, with a Kd = 4.3 · 10^-7^ / 5.9 · 10^-8^ M, while the affinity of BtuG1 toward Cbi sits at Kd = 5.8 · 10^-5^ / 1.4 · 10^-5^ M; an approximately 100-fold difference. These differences in Cbi affinities confirm BtuG homologs to possess different specificities toward the precursor Cbi, and reinforce the results carried out for cobalamin competition studies, that found the *locus*2, composed by BtuG2 and BtuB2, as the most efficient for cobalamin transport (Degnan et al., 2014). On the other hand, the three BtuG homologs bear equally high affinity for the two Cbl variants tested, CNCbl and OHCbl in the range of 1-10 pM, and, therefore, we cannot highlight a specific homolog contributing the most to the organism’s fitness toward B12 transport. In previous reports, the measured affinity of BtuG2 toward both Cbl and Cbi was ∼1.9 · 10^-13^ M (Wexler et al., 2018). That is, approximately a 7-orders-of-magnitude affinity increase for Cbi and 10-times higher affinity for Cbl than measured in our study. We believe the absolute affinity values of BtuG2 measured using GCI technology to be of higher accuracy than surface plasmon resonance (SPR), since as per the SPR manufacturer specification (Biacore T100, GE Healthcare), these affinities are outside the trust range of the SPR device.

Based on the high-resolution structures of molecular interactions between all BtuG homologs and their substrates, we aimed at determining which residues were important for affinity increase and decrease. Interestingly, Trp272 in BtuG2, which rotates upon substrate binding (Fig. 3c), makes a hydrophobic interaction with Cbl in the same position. Ala261 is present at the equivalent position in BtuG3, which does not provide any contact with the substrate (Fig. 3b and c). The measured affinity for Cbi in BtuG2 is Kd = 4.3 · 10^-7^ / 5.9 · 10^-8^ M, while BtuG3 can bind Cbi with a Kd = 3.1 · 10^-5^ / 3.9 · 10^-6^ M, a 100-fold difference in affinity. We designed a BtuG3 A261W mutant that aimed to supplement this missing interaction at the binding site of BtuG3, therefore, increasing the affinity of BtuG3 toward Cbi. These efforts were partially successful as the affinity for Cbi on the BtuG3 A261W mutant was 10-fold higher (in one component of the heterologous binding), with Kd = 4.3 · 10^-6^ / 3.3 · 10^-6^ M. Inversely, removing the Trp272 interaction from BtuG2 in a W272A mutant decreases the affinity 10/20-fold in both populations, to a Kd = 5.0 · 10^-6^ / 2.0 · 10^-6^ M. Therefore, we can conclude that the hydrophobic interaction provided by Trp272 directly influences the stability of the interaction to Cbi, and hence, its affinity for the substrate, as the indolic sidechain can pivot to accommodate the corrin ring. This finding is consistent with the recent structural study on BtuG2 bound to an AdoCbl, where Trp272 sidechain rotates outwards to accommodate the bulky aldehyde upper ligand of AdoCbl (Abellon-Ruiz et al., 2023). Tryptophan has also been reported to perform this adaptive function for Cbl binding on the ABC transporter substrate-binding subunit BtuF, where a tryptophan residue was key to stabilize the interaction with Cbi (Mireku et al., 2017).

Overall, these results establish the BtuG family as super-binders for cobalamin, with extreme affinity levels, comparable to those of proteins with medical and biotechnological applications, e.g., Streptavidin and Nanobodies; and set the ground for further research on targeted drug delivery to human gut microbiota and affinity-based essays using Cbl-BtuG complex as a vector for protein detection and purification.

## Material and Methods

### BtuG homologs cloning

Synthetic DNA sequences (GenScript) containing the sequences of the globular domains (excluding N-terminal proteolipidic signal peptide) of BtuG1-2-3, codon optimized for *E. coli* (Suppl. Table 1) were cloned into a pBAD24 with ampicillin resistance using the USER cloning method (New England Biolabs, Table 2). All sequences were cloned including a 10x histidine tag located in the N-terminus for BtuG1-3 and the C-terminus for BtuG2 and all its mutants. Between the tag and the protein sequence a 3C cleavage sequence was included, to allow subsequent cleavage of the tag during the purification (All primers are listed on Suppl. Table 1).

Each construct is transformed into a chemically competent MC1061 strain of *E. coli*. A colony PCR using Taq Polymerase (New England Biolabs) on positive resistant colonies can help identifying false positives for Ampicillin resistance. Positive strains are checked for mutations using DNA sequencing (Eurofins).

### BtuG homologs expression

All BtuG constructs were expressed and grown in the same conditions, obtaining similar expression levels. A pre-saturated LB culture of *E. coli* bearing the corresponding BtuG vector inoculated large 5 L flasks containing 2 L LB medium at a 1/100 v:v. Cells were shaken at 37 °C until OD at 600 nm reached 0.6-0.8. Then protein overexpression was induced using 0.01% L-Arabinose and left shaking at 37 °C for 3 hours. Then, cells were harvested, washed with 50 mM K-P_i_ pH 7.5, glycerol 10%, flash frozen in liquid nitrogen and stored at -80 °C. Cells were disrupted with a Maximator Systems at 20 kPsi and 4 °C and supplemented with 200 μM PMSF, 1 mM MgSO_4_ and DNase I. Cell debris and membrane lipids were removed by ultracentrifugation for 30 min at 158,420×*g* (average) and 4 °C.

### BtuG homologs purification

The supernatant is incubated with 0.25 ml of Superflow Ni^2+^-NTA Sepharose per L of LB culture equilibrated with 20 column volumes (CV) of wash buffer (20 mM Tris/HCl pH 7.5, 100 mM NaCl, 50 mM imidazole/HCl pH 7.5). Unbound protein was allowed to flow through and the column was washed with 40 CV wash buffer. For crystallography experiments, the engineered histidine tag was cleaved using 3C protease. For this, the column resin was resuspended adding 0.1 ml per L of LB of 3C protease in purification buffer containing 1.2 mg of 3C and 50 mM Tris-HCl pH 8.0, 150 mM NaCl, 20% glycerol, 10 mM EDTA and 1 mM DTT. The final cleavage solution was adjusted to 100 mM imidazole pH 8.0 for an optimal cleavage. The reaction was incubated for 2 hours at 4 °C with slow rocking. The cleaved protein was then allowed to flow through using wash buffer. The 3C protease used for this experiment contains an engineered histidine-tag, remaining bound to the column after the cleavage. For kinetic assays, the protein was eluted off the resin using wash buffer adjusted to 500 mM imidazole/HCl pH 7.5. Both tagged and cleaved BtuG samples were then loaded on a NAP-10 desalting column (kinetic assays, GE healthcare) for buffer exchange to 20 mM Tris/HCl pH 7.5 and 100 mM NaCl. All further experiments were performed with this buffer composition, unless otherwise stated.

### Crystallization, data collection and structure determination

For each crystallization trial, following purification, the protein was concentrated and a stoichiometric excess of substrate, either Cbi or CNCbl, was added. Each of the proteins were mixed with the crystallization buffer in a 2:1 ratio, respectively. All crystals were obtained using sitting drop vapor diffusion via commercial or custom screening kits. Following concentration to 28 mg/mL and substrate binding, BtuG1 was screened for crystals using the MCSG1 crystallization kit (Anatrace Products). Several hits were observed after 24 hours at 4℃, the most promising of which were further optimised to yield X-ray quality diffracting crystals using the condition: 0.1 M Bis-Tris propane HCl, pH 7.0, 1.4 M sodium malonate, pH 7.0. BtuG2 was concentrated to 22 mg/mL prior to substrate addition and screened for crystals using the MCSG3 crystallization kit (Anatrace Products). Several hits were observed after 48 hours at 4 ℃ using the following condition: 1 M ammonium sulfate, 0.1 M HEPES NaOH, pH 7.0, 0.5 % (w/v) PEG 8000. BtuG3 was concentrated to 43 mg/mL, an excess of Cbi added, crystal conditions screened using the MCSG2 crystallization kit (Anatrace Products). X-ray quality crystals grew after 24 hours at 4 ℃ using the condition: 0.2 M potassium nitrate, pH 6.9, 20 % (w/v) PEG 3350. All crystals were briefly soaked in 30 % (v/v) glycerol for cryo-protection and subsequently flash-frozen in liquid nitrogen in preparation for diffraction experiments. Data were collected at beam line ID23-1 (ESRF, Grenoble). Crystals of the BtuG series diffracted between 1.5 and 2.0 Å resolution (Table 1). Data were processed with XDS (Kabsch, 2010). BtuG1 was solved by Molecular Replacement with Phaser (McCoy et al., 2007) using an AlphaFold (Jumper et al., 2021) predicted model from the given sequence. BtuG2 was solved using an automated MoRDa (Vagin & Lebedev, 2015) Molecular Replacement workflow. BtuG3 was solved by Molecular Replacement with Phaser, using the model of BtuG2. Manual rebuilding was performed with COOT (Emsley et al., 2010) and refinement with Refmac (Murshudov et al., 2011). The refined models and associated structure factors were deposited into the PDB repository with the following IDs: 8BAI, 8BAZ, 8BB0, 8PTD, 8PTF, 8PUJ. All structural figures were prepared with an open-source version of PyMOL (https://sourceforge.net/projects/pymol/). ButG1-Cbi crystals experienced severe anisotropy (hence low completeness), and all attempts to improve the quality of crystals were unsuccessful.

**Table 1:** Data collection and refinement statistics (molecular replacement).

|  | **BtuG1-Cbi** | **BtuG1-CNCbl** | **BtuG2-Cbi** | **BtuG2-OHCbl** | **BtuG2-CNCbl** | **BtuG3-Cbi** |
| --- | --- | --- | --- | --- | --- | --- |
| **PDB-ID** | **8PTF** | **8PTD** | **8BAZ** | **8BB0** | **8BAI** | **8PUJ** |
| Wavelength | 0.8731 | 0.8731 | 1.0 | 0.9677 | 1.0 | 0.8731 |
| Resolution range | 45.43 - 1.974<br>(2.045 - 1.974) | 34.26 - 1.58<br>(1.636 - 1.58) | 44.16 - 2.0<br>(2.071 - 2.0) | 36.85 - 1.5<br>(1.554 - 1.5) | 36.88 - 1.692<br>(1.753 - 1.692) | 44.02 - 1.663<br>(1.722 - 1.663) |
| Space group | C 2 2 21 | C 2 2 21 | P 1 21 1 | P 1 21 1 | P 1 | P 21 21 2 |
| *a*, *b*, *c* (Å) | 52.724<br>145.391<br>116.382 | 52.471<br>147.665<br>116.043 | 49.177<br>100.593<br>79.579 | 49.110<br>101.10<br>79.893 | 49.24<br>79.936<br>101.047 | 107.993<br>151.985<br>45.618 |
| $\alpha$, $\beta$, $\gamma$ (°) | 90 90 90 | 90 90 90 | 90 97.6 90 | 90 97.81 90 | 89.966<br>89.892<br>82.277 | 90 90 90 |
| Unique reflections | 18162<br>(150) | 62037<br>(6158) | 51786<br>(5152) | 123234 | 163972<br>(15974) | 88345<br>(8238) |
| Completeness (%) | 56.89<br>(4.76) | 99.86<br>(99.87) | 99.84<br>(99.90) | 95.64 (97.44) | 96.24<br>(93.53) | 98.94<br>(93.73) |
| Mean I/sigma(I) | 12.33 | 14.70 | 9.54 | 14.05 | 10.14 | 10.90 |
| Wilson B-factor | 26.73 | 19.63 | 27.09 | 11.37 | 15.82 | 16.17 |
| R-meas | 0.10 | 0.06 | 0.06 | 0.08 | 0.08 | 0.08 |
| CC<sub>1/2</sub> | 91.3<br>(27.0) | 98.2 (28.1) | 98.5<br>(32.3) | 93.0 (30.2) | 99.1 (27.7) | 92.2 (29.0) |
| Reflections used in refinement | 18158<br>(150) | 62035<br>(6158) | 51757<br>(5149) | 117831<br>(11971) | 163912<br>(15936) | 88343<br>(8238) |
| Reflections used for R-free | 4216<br>(419) | 3096 (315) | 2530<br>(231) | 5871 (600) | 1978 (195) | 4312 (413) |
| R-work | 0.1835<br>(0.2532) | 0.1909<br>(0.2648) | 0.1854<br>(0.2087) | 0.1565<br>(0.2310) | 0.1457<br>(0.2263) | 0.1825<br>(0.2957) |
| R-free | 0.2002<br>(0.3229) | 0.2102<br>(0.3099) | 0.2512<br>(0.2980) | 0.1871<br>(0.2770) | 0.1816<br>(0.2582) | 0.2111<br>(0.3263) |
| Number of non-hydrogen atoms | 2817 | 3045 | 5882 | 6137 | 12018 | 5879 |
| macromolecules | 2668 | 2662 | 5247 | 5269 | 10494 | 5257 |
| ligands | 78 | 131 | 397 | 360 | 545 | 85 |
| solvent | 71 | 252 | 238 | 508 | 997 | 537 |
| Protein residues | 342 | 342 | 641 | 641 | 1282 | 644 |
| RMS(bonds) | 0.009 | 0.016 | 0.011 | 0.017 | 0.017 | 0.013 |
| RMS(angles) | 1.76 | 2.27 | 1.69 | 2.11 | 2.09 | 2.11 |
| Ramachandran favored (%) | 95.29 | 96.76 | 94.19 | 94.03 | 94.19 | 94.84 |
| Ramachandran allowed (%) | 4.41 | 2.94 | 5.02 | 5.02 | 5.02 | 4.22 |
| Ramachandran outliers (%) | 0.29 | 0.29 | 0.78 | 0.94 | 0.78 | 0.94 |
| Rotamer outliers (%) | 1.01 | 1.69 | 1.41 | 0.87 | 0.97 | 0.17 |
| Clashscore | 7.57 | 7.62 | 9.60 | 7.16 | 7.46 | 6.20 |
| Average B-factor | 21.00 | 24.48 | 32.09 | 18.32 | 20.50 | 21.04 |
| macromolecules | 20.19 | 23.35 | 31.29 | 16.08 | 18.76 | 19.84 |
| ligands | 49.95 | 26.87 | 37.58 | 27.34 | 24.24 | 37.42 |
| solvent | 19.50 | 35.21 | 40.62 | 35.12 | 36.79 | 30.22 |

\*Statistics for the highest resolution shell are shown in parentheses.

**Table 2.** Affinity constants of each BtuG homolog and mutant toward CNCbl, OHCbl and Cbi. Average k_on_, k_off_ and dissociation constant (Kd) for every substrate affinity experiment, measured on each BtuG construct, using GCI technology. Between parenthesis the number of experiment replicates (See methods). Error represents the standard deviation for that affinity value.

| Construct | Substrate | Association ($k_a$, $M^{-1}s^{-1}$) | Dissociation ($k_d$, $s^{-1}$) | Equilibrium ($K_d$, M) |
| --- | --- | --- | --- | --- |
| **BtuG1** | CNCbl | $8.3 \cdot 10^7 \pm 1.1 \cdot 10^{-7}$ (2) | $1.1 \cdot 10^{-4} \pm 0.3 \cdot 10^{-4}$ (2) | $1.3 \cdot 10^{-12} \pm 0.4 \cdot 10^{-12}$ (2) |
| | OHCbl | $5.1 \cdot 10^7$ (1) | $1.6 \cdot 10^{-4}$ (1) | $3.0 \cdot 10^{-12}$ (1) |
| | Cbi | $9.9 \cdot 10^2 \pm 1.8 \cdot 10^2$ / $3.1 \cdot 10^2 \pm 0.0$ (2) | $6.0 \cdot 10^{-2} \pm 3.1 \cdot 10^{-2}$ / $3.9 \cdot 10^{-3} \pm 1.2 \cdot 10^{-3}$ (2) | $5.8 \cdot 10^{-5} \pm 2.2 \cdot 10^{-5}$ / $1.4 \cdot 10^{-5} \pm 0.5 \cdot 10^{-5}$ (2) |
| **BtuG1 · 10180W** | CNCbl | $3.2 \cdot 10^7$ (1) | $1.3 \cdot 10^{-2}$ (1) | $4.2 \cdot 10^{-10}$ (1) |
| | OHCbl | $1.8 \cdot 10^6$ (1) | $1.5 \cdot 10^{-2}$ (1) | $0.9 \cdot 10^{-8}$ (1) |
| | Cbi | $6.4 \cdot 10^2 \pm 0.4 \cdot 10^2$ / $7.3 \cdot 10^2 \pm 1.8 \cdot 10^2$ (2) | $4.7 \cdot 10^{-2} \pm 0.2 \cdot 10^{-2}$ / $3.6 \cdot 10^{-3} \pm 1.1 \cdot 10^{-3}$ (2) | $1.2 \cdot 10^{-4} \pm 0.6 \cdot 10^{-4}$ / $6.9 \cdot 10^{-5} \pm 0.2 \cdot 10^{-5}$ (2) |
| **BtuG1 R244W** | CNCbl | $2.5 \cdot 10^7$ (1) | $4.5 \cdot 10^{-3}$ (1) | $1.8 \cdot 10^{-10}$ (1) |
| | OHCbl | $1.2 \cdot 10^6$ (1) | $3.7 \cdot 10^{-2}$ (1) | $3.2 \cdot 10^{-8} / 2.6 \cdot 10^{-11}$ (1) |
| | Cbi | $4.2 \cdot 10^2 \pm 0.9 \cdot 10^2$ / $3.7 \cdot 10^2 \pm 0.7 \cdot 10^2$ (2) | $3.4 \cdot 10^{-2} \pm 1.0 \cdot 10^{-2}$ / $2.5 \cdot 10^{-3} \pm 0.7 \cdot 10^{-3}$ (2) | $8.8 \cdot 10^{-5} \pm 5.9 \cdot 10^{-5}$ / $6.9 \cdot 10^{-6} \pm 1.1 \cdot 10^{-6}$ (2) |
| **BtuG1 Y290W** | CNCbl | $1.9 \cdot 10^8$ (1) | $1.0 \cdot 10^{-7}$ (1) | $5.2 \cdot 10^{-16}$ (1) |
| | OHCbl | $2.6 \cdot 10^7$ (1) | $3.7 \cdot 10^{-6}$ (1) | $1.4 \cdot 10^{-13}$ (1) |
| | Cbi | $9.2 \cdot 10^2 \pm 1.2 \cdot 10^2$ / $2.3 \cdot 10^2 \pm 1.1 \cdot 10^2$ (3) | $3.2 \cdot 10^{-2} \pm 1.3 \cdot 10^{-2}$ / $1.7 \cdot 10^{-3} \pm 0.5 \cdot 10^{-3}$ (3) | $3.6 \cdot 10^{-5} \pm 1.7 \cdot 10^{-5}$ / $8.5 \cdot 10^{-6} \pm 3.4 \cdot 10^{-6}$ (3) |
| **BtuG2** | CNCbl | $1.4 \cdot 10^8 \pm 0.8 \cdot 10^8$ (3) | $1.5 \cdot 10^{-4} \pm 1.1 \cdot 10^{-4}$ (3) | $1.7 \cdot 10^{-12} \pm 1.4 \cdot 10^{-12}$ (3) |
| | OHCbl | $2.6 \cdot 10^7$ (1) | $5.5 \cdot 10^{-5}$ (1) | $2.2 \cdot 10^{-12}$ (1) |
| | Cbi | $4.5 \cdot 10^4 \pm 2.5 \cdot 10^4 /$<br>$1.4 \cdot 10^4 \pm 0.2 \cdot 10^4$ (4) | $1.5 \cdot 10^{-2} \pm 0.5 \cdot 10^{-2} /$<br>$7.8 \cdot 10^{-4} \pm 4.2 \cdot 10^{-4}$ (4) | $4.3 \cdot 10^{-7} \pm 2.3 \cdot 10^{-7} /$<br>$5.9 \cdot 10^{-8} \pm 2.9 \cdot 10^{-8}$ (4) |
| **BtuG2 · 1055A** | CNCbl | $2.6 \cdot 10^7$ (1) | $4.0 \cdot 10^{-5}$ (1) | $1.5 \cdot 10^{-12}$ (1) |
|  | OHCbl | - | - | - |
| | Cbi | $1.0 \cdot 10^5 / 1.8 \cdot 10^4$ (1) | $1.3 \cdot 10^{-2} / 6.5 \cdot 10^{-4}$ (1) | $1.2 \cdot 10^{-7} / 3.6 \cdot 10^{-8}$ (1) |
| **BtuG2 F58A** | CNCbl | $2.4 \cdot 10^8$ (1) | $2.4 \cdot 10^{-4}$ (1) | $1.0 \cdot 10^{-12}$ (1) |
|  | OHCbl | - | - | - |
| | Cbi | $6.3 \cdot 10^3 \pm 4.2 \cdot 10^3 /$<br>$4.5 \cdot 10^3 \pm 4.1 \cdot 10^3$ (3) | $7.0 \cdot 10^{-3} \pm 1.9 \cdot 10^{-3} /$<br>$4.0 \cdot 10^{-5} \pm 5.7 \cdot 10^{-5}$ (3) | $3.5 \cdot 10^{-6} \pm 4.9 \cdot 10^{-6} /$<br>$1.9 \cdot 10^{-8} \pm 3.2 \cdot 10^{-8}$ (3) |
| **BtuG2 Q93A** | CNCbl | $1.5 \cdot 10^7$ (1) | $6.0 \cdot 10^{-7}$ (1) | $4.0 \cdot 10^{-14}$ (1) |
|  | OHCbl | - | - | - |
| | Cbi | $2.0 \cdot 10^3 / 1.0 \cdot 10^2$ (1) | $9.8 \cdot 10^{-3} / 1.7 \cdot 10^{-4}$ (1) | $5.0 \cdot 10^{-6} / 1.7 \cdot 10^{-6}$ (1) |
| **BtuG2 Q219A** | CNCbl | $4.9 \cdot 10^7$ (1) | $2.0 \cdot 10^{-5}$ (1) | $4.1 \cdot 10^{-13}$ (1) |
|  | OHCbl | - | - | - |
| | Cbi | $3.2 \cdot 10^3 \pm 1.4 \cdot 10^3 /$<br>$2.4 \cdot 10^2 \pm 2.2 \cdot 10^2$ (2) | $5.6 \cdot 10^{-3} \pm 3.0 \cdot 10^{-3} /$<br>$7.6 \cdot 10^{-6} \pm 7.5 \cdot 10^{-6}$ (2) | $1.7 \cdot 10^{-6} \pm 0.4 \cdot 10^{-6} /$<br>$1.8 \cdot 10^{-8} \pm 1.9 \cdot 10^{-8}$ (2) |
| **BtuG2 W272A** | CNCbl | $5.6 \cdot 10^7$ (1) | $1.9 \cdot 10^{-6}$ (1) | $3.5 \cdot 10^{-14}$ (1) |
|  | OHCbl | - | - | - |
| | Cbi | $2.3 \cdot 10^3 \pm 0.4 \cdot 10^3 /$<br>$3.9 \cdot 10^2 \pm 0.6 \cdot 10^2$ (3) | $1.1 \cdot 10^{-2} \pm 0.3 \cdot 10^{-2} /$<br>$9.1 \cdot 10^{-4} \pm 8.9 \cdot 10^{-4}$ (2) | $5.0 \cdot 10^{-6} \pm 0.1 \cdot 10^{-6} /$<br>$2.0 \cdot 10^{-6} \pm 2.7 \cdot 10^{-6}$ (2) |
| **BtuG3** | CNCbl | $3.8 \cdot 10^7 \pm 1.5 \cdot 10^7$ (3) | $8.6 \cdot 10^{-4} \pm 7.7 \cdot 10^{-4}$ (3) | $1.7 \cdot 10^{-11} \pm 1.5 \cdot 10^{-11}$ (3) |
| | OHCbl | $2.8 \cdot 10^8 \pm 2.6 \cdot 10^8$ (2) | $7.4 \cdot 10^{-4} \pm 4.7 \cdot 10^{-4}$ (2) | $2.8 \cdot 10^{-11} \pm 3.9 \cdot 10^{-11}$ (2) |
| | Cbi | $2.0 \cdot 10^3 \pm 2.2 \cdot 10^3 /$<br>$4.2 \cdot 10^2 \pm 0.6 \cdot 10^2$ (3) | $2.4 \cdot 10^{-2} \pm 0.3 \cdot 10^{-2} /$<br>$1.7 \cdot 10^{-3} \pm 0.4 \cdot 10^{-3}$ (3) | $3.1 \cdot 10^{-5} \pm 2.3 \cdot 10^{-5} /$<br>$3.9 \cdot 10^{-6} \pm 1.0 \cdot 10^{-6}$ (3) |
| **BtuG3 A261W** | CNCbl | $7.8 \cdot 10^7$ (1) | $2.3 \cdot 10^{-3}$ (1) | $3.0 \cdot 10^{-11}$ (1) |
| | OHCbl | $3.6 \cdot 10^6$ (1) | $5.1 \cdot 10^{-4}$ (1) | $1.4 \cdot 10^{-10}$ (1) |
| | Cbi | $5.7 \cdot 10^3 \pm 0.2 \cdot 10^3 /$<br>$6.2 \cdot 10^2 \pm 0.9 \cdot 10^2$ (2) | $2.5 \cdot 10^{-2} \pm 0.1 \cdot 10^{-2} /$<br>$2.0 \cdot 10^{-3} \pm 0.1 \cdot 10^{-3}$ (2) | $4.3 \cdot 10^{-6} \pm 0.0 /$<br>$3.3 \cdot 10^{-6} \pm 0.3 \cdot 10^{-6}$ (2) |

### Kinetic measurements

Binding kinetics for BtuG samples were measured in a WAVE System microfluidics device (Creoptix, AG). A fresh PCH-NTA microfluidics chip with a nickel coating was conditioned with borate buffer (80 µl borate 50 mM pH 9.0, NaCl 1 M), cleaned with 80 µl EDTA/NaOH 350 mM pH 8.0 and pre-equilibrated with running buffer (20 mM Tris/HCl pH 7.5, 150 mM NaCl) until the signal became stable. Between solutions, the chip was always rinsed with running buffer. During binding kinetics experiments, one out of the four channels present in PCH-NTA chips was left empty of sample protein, and another channel was bound with 3C protease for referencing. Two channels were then left for BtuG sample loading. Every kinetic cycle, each channel was cleaned from nickel ions with EDTA/NaOH 350 mM pH 8.0, activated for affinity binding with NiSO_4_ 0.5 mM, loaded with purified BtuG sample including the engineered histidine tag in solutions ranging from 0.5-to-1 mg/ml until 125 pg/mm^2^ for Cbl measurements, and 2000 pg/mm^2^ for Cbi. After BtuG sample loading, the chip was washed with running buffer until the signal was stable. Then, substrate solution (running buffer supplemented with the appropriate concentration of substrate) was loaded onto the chip. For double referencing, frequent blanks were loaded where running buffer substitutes the substrate buffer during the kinetic cycle. Every experiment consisted of at least five different substrate concentrations, each of them adjusted to cover the range where the lowest substrate concentration shows negligible binding, and the highest concentration reaches saturation (Suppl. Fig. 4). Data was analyzed using WAVE modelling software. All Cbl models apply a mass transport limitation calculation due to extremely high association constant of BtuG homologs. All Cbi models apply a heterologous ligand calculation due to the biphasic nature of Cbi binding.

### Mass spectrometry

BtuG1 was purified for exact mass determination as described above and concentrated to approximately 2 mg/mL for mass spectrometry analysis. BtuG1 protein was diluted in a 1:10 (v/v) ratio with 0.1% formic acid and 10 μL were injected into an UFLC-XR LC system (Shimadzu), connected to a Q-Exactive mass spectrometer (Thermo Fisher Scientific) and separated on a 2.1 mm × 50 mm Acquity UPLC BEH C18, 1.7 μm (Waters). Solvent A was H_2_O with 0.1% formic acid and solvent B was acetonitrile with 0.1% formic acid. The following mobile phase gradient was delivered at a flow rate of 0.6 ml min^−1^ starting with a mixture of 30 % solvent B. Solvent B was increased to 90 % over 6 minutes with a linear gradient and kept at this concentration for 5 minutes. Solvent B was reduced to 30 % in 0.1 minutes and kept for 3.9 minutes resulting in a total elution time of 15 minutes. The column temperature was kept constant at 40 °C. The mass spectrometer was operated in positive mode. Full scan MS spectra were acquired for 10 minutes from *m/z* 1000 to 2000 at a target value of 1 × 106 and a max IT of 500 ms with a resolution of 70,000 at m/z 200. Scans were resolved with Thermo fisher BioPharma Finder software.

## Supplementary information

**Supplementary table 1.**
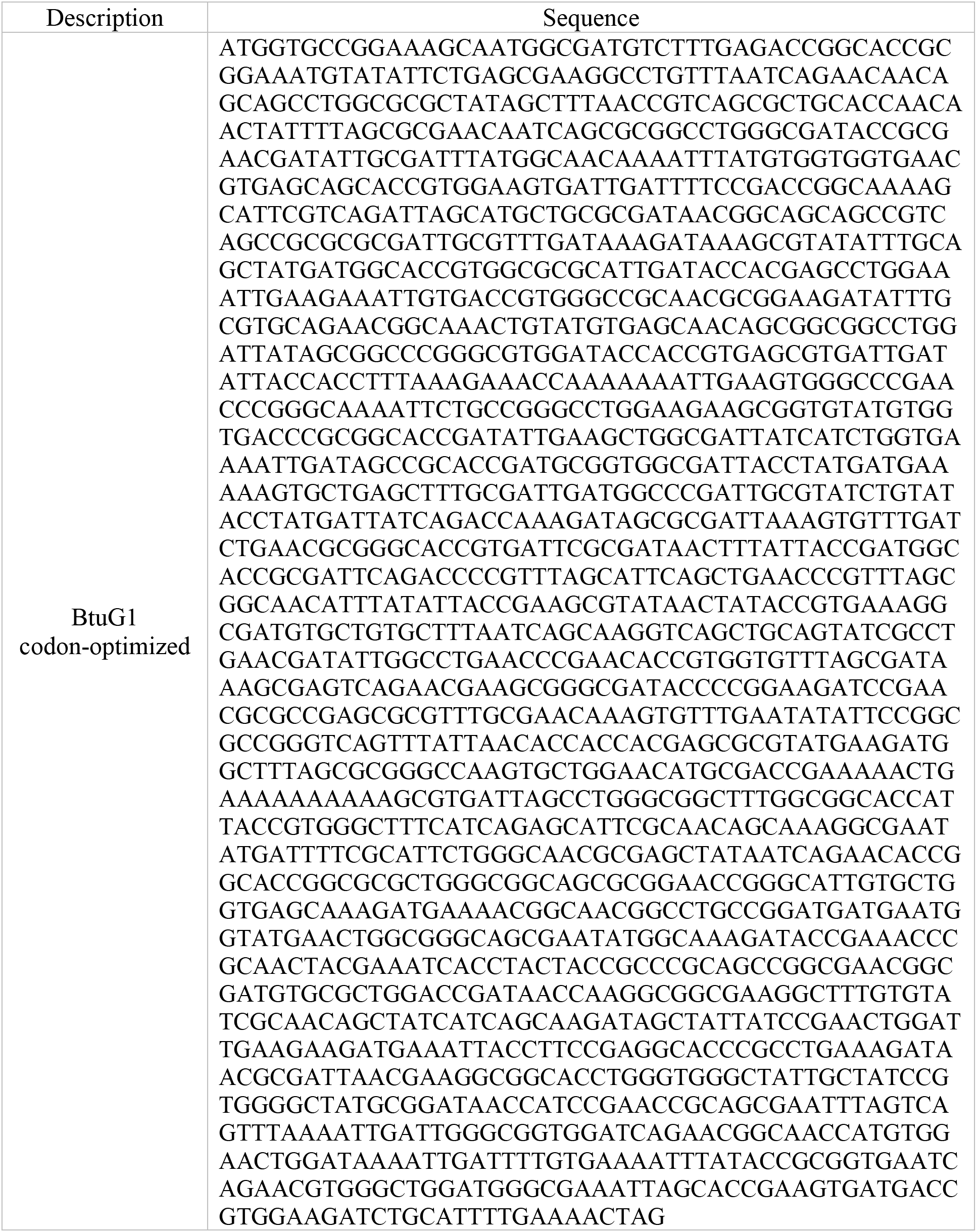
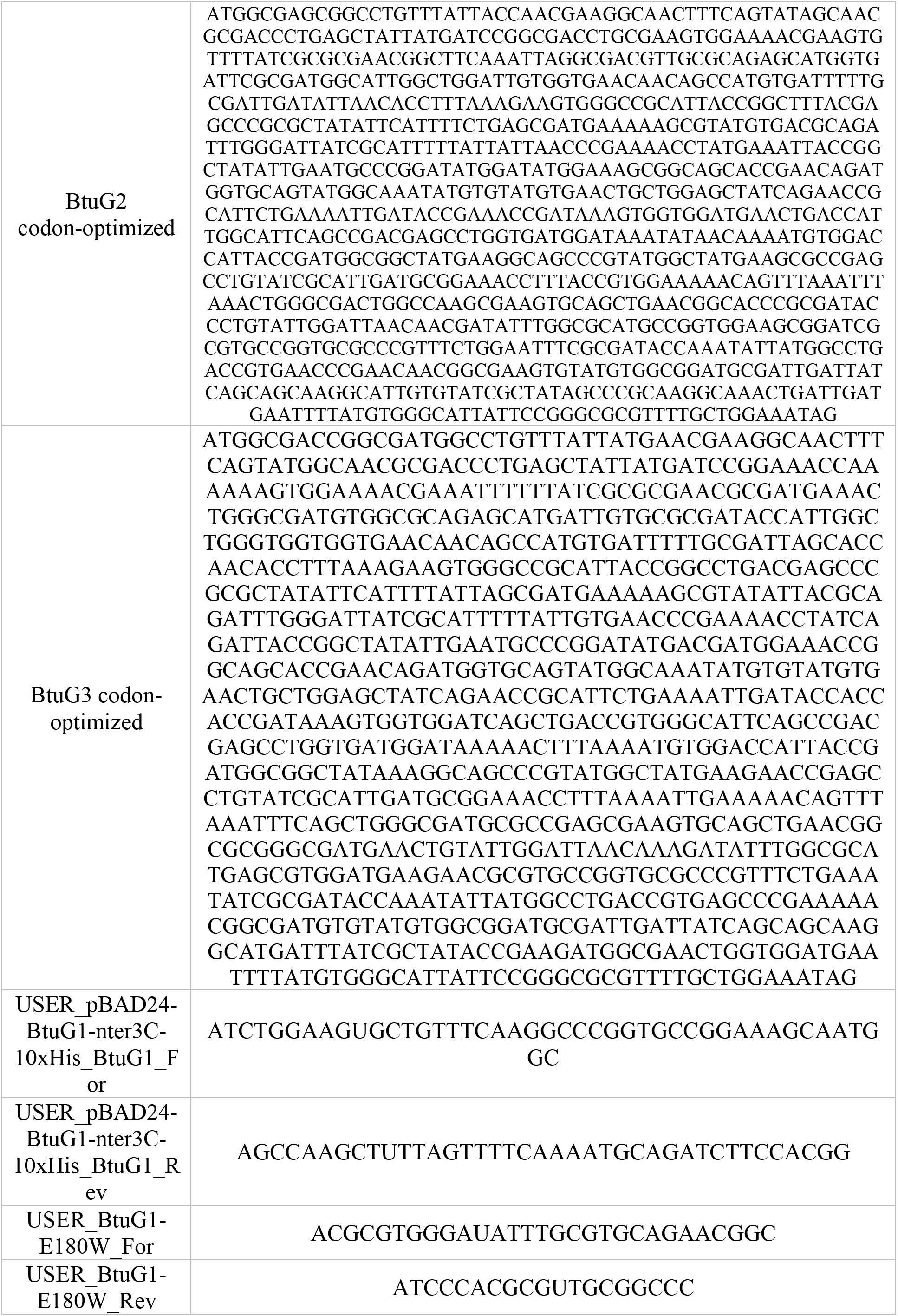
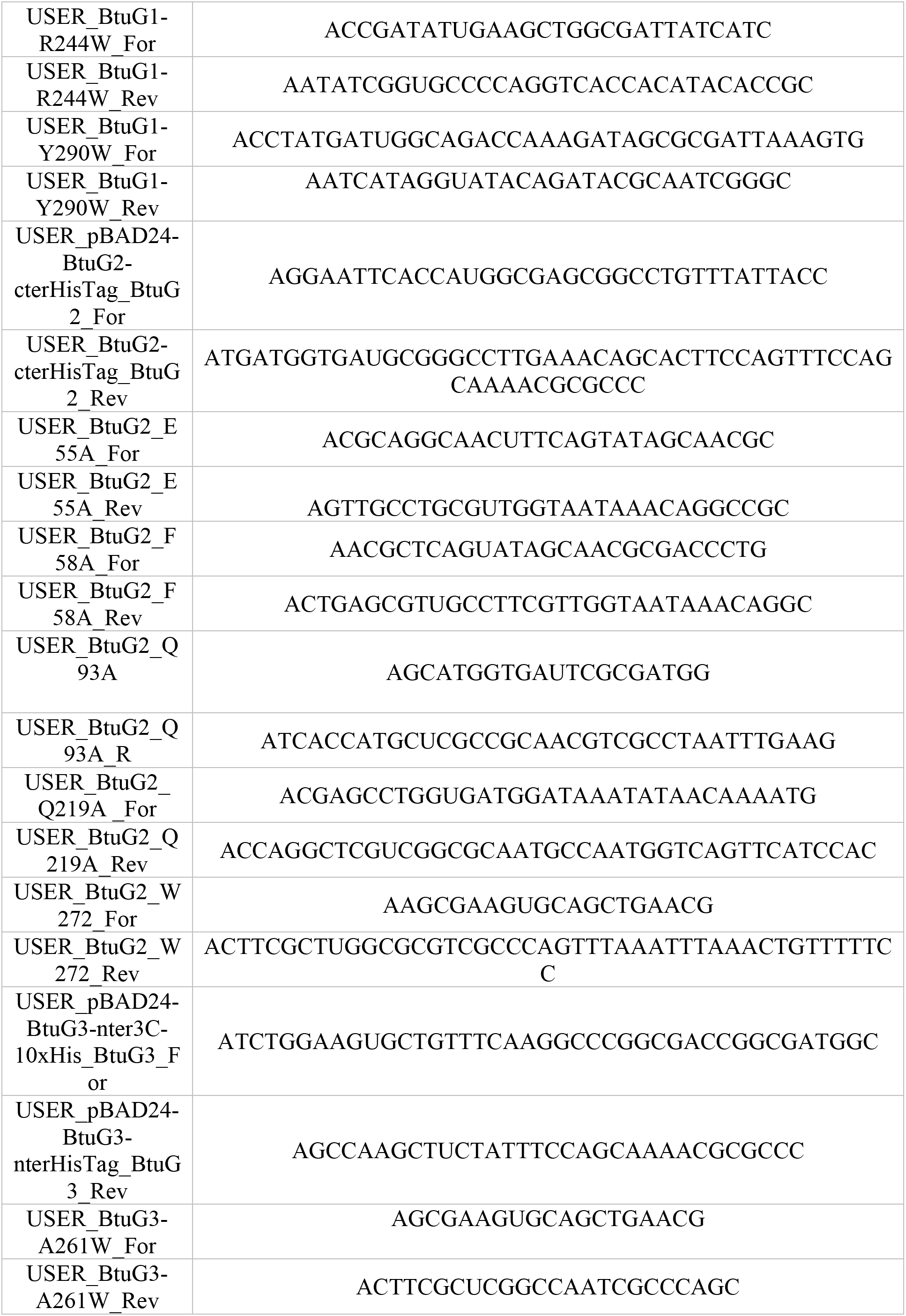
Codon-optimized DNA sequences of BtuG homologs and primers used for this study.

**Supplementary figure 1.**
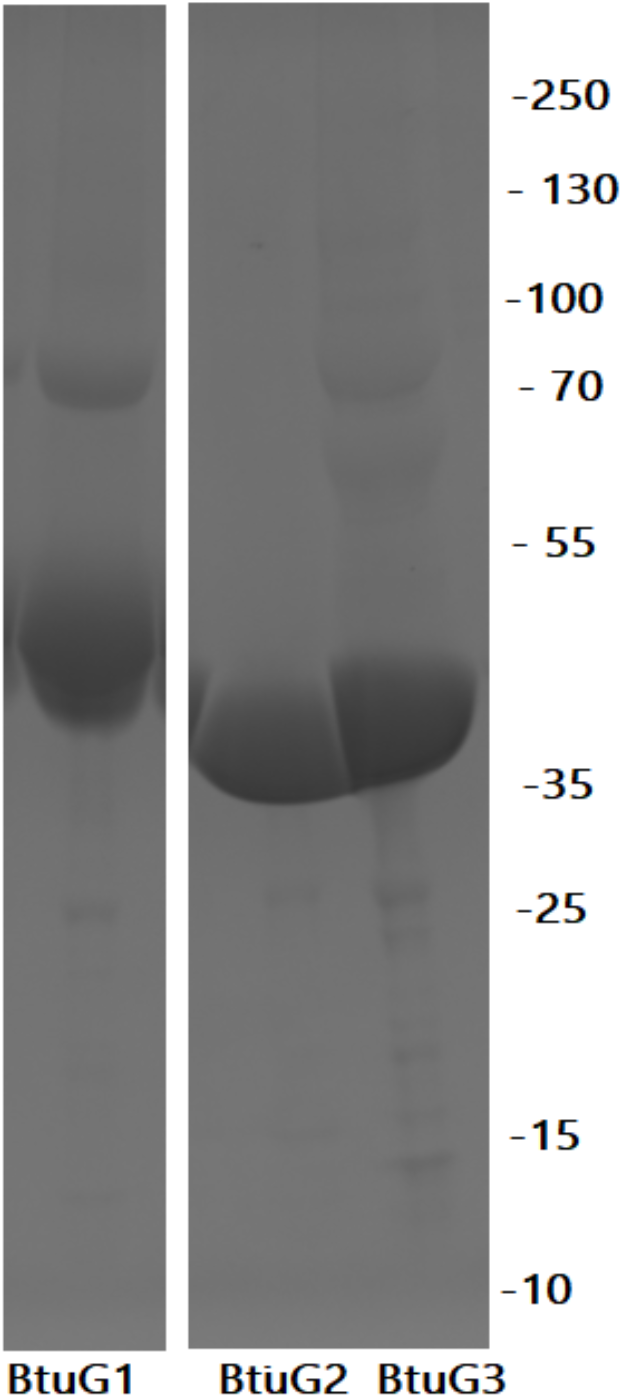
SDS-PAGE gel of purified BtuG1-2-3 homologs. Every lane is labelled for the homolog construct loaded, on the right, the molecular weight in kDa.

**Supplementary figure 2.**
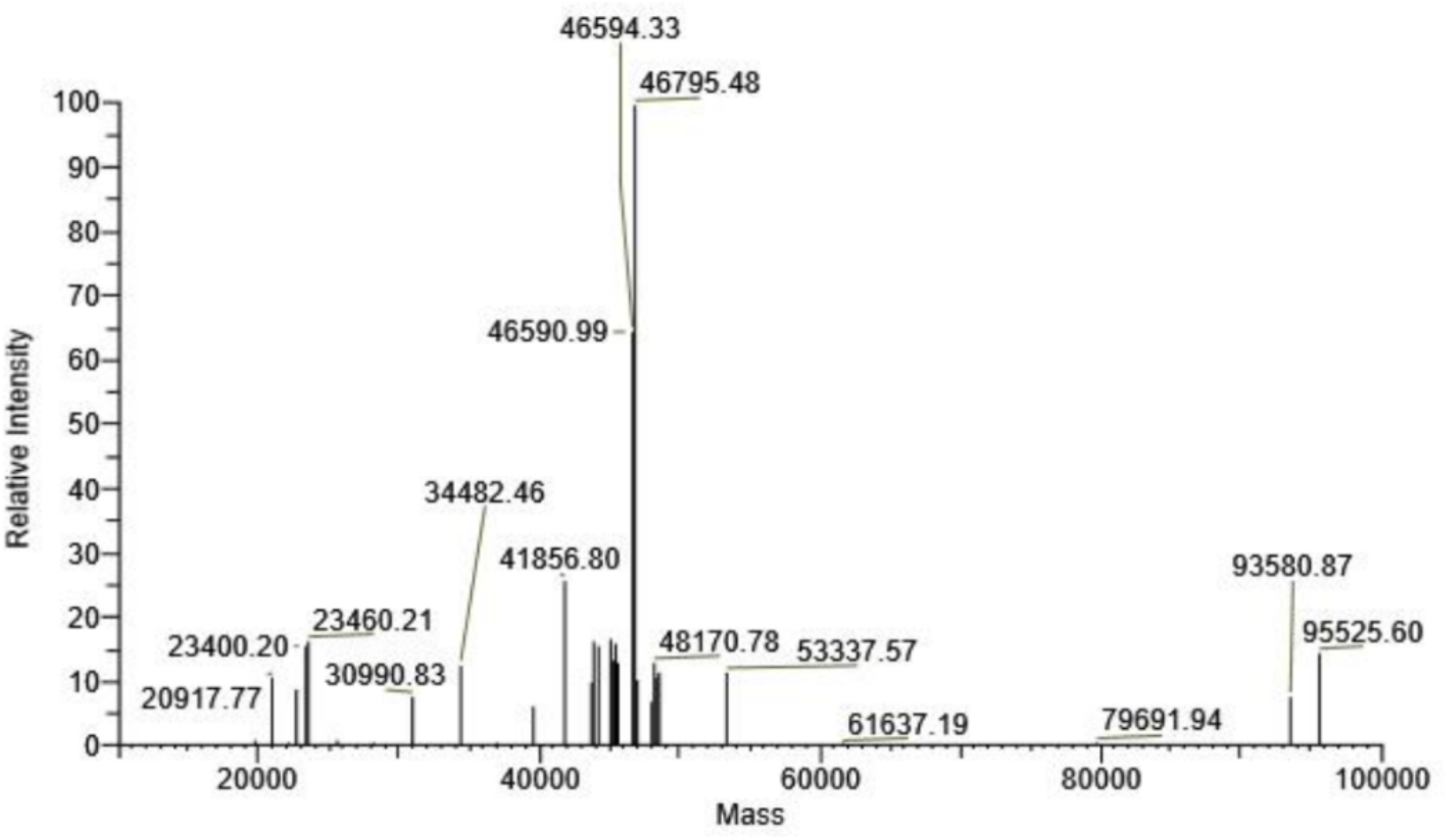
Purified BtuG1 (expected mass of 69607 Da) cleaved using 3C protease yields a prominent peak at 46795 Da, corresponding to a truncated form with loss of a 22812 Da C terminal peptide. The potential cleavage site of the truncated version is between L442 and G443.

**Supplementary figure 3.**
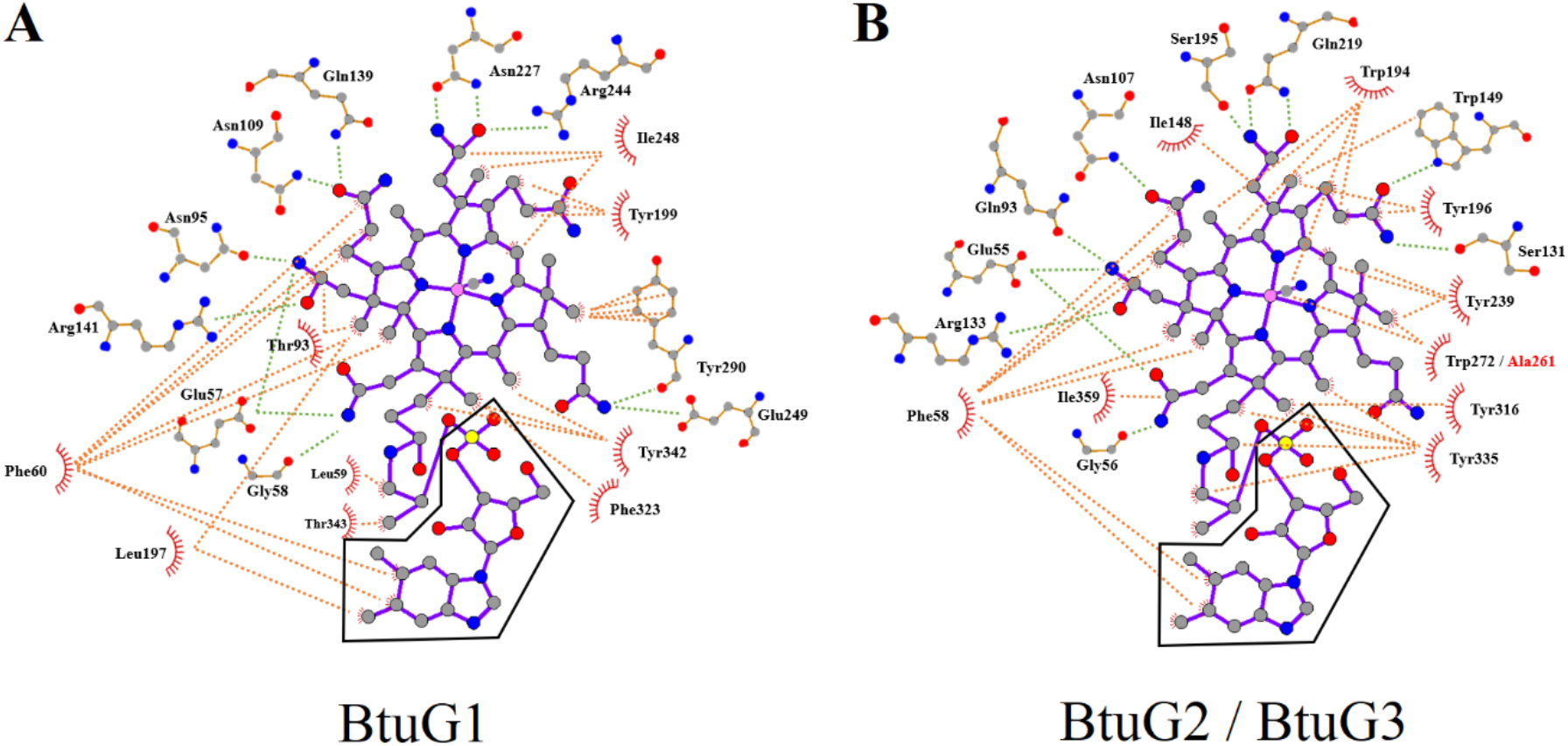
2D interaction diagram. Protein-ligand interactions at the binding site of BtuG1 (A) and BtuG2 and BtuG3 (B) toward CNCbl (coloured in purple) and Cbi (excludes the DMB moiety inside the black frame). Polar interactions (polar atoms closer than 3.5 Å) are shown in green dashed lines, and hydrophilic interactions (non-polar atoms closer than 4.0 Å) in orange. Residues involved in hydrophilic interactions are summarized as a red crown pointing toward their interaction site. Carbon atoms are represented by grey (residues) or purple (ligand) circles, nitrogen in blue, oxygen in red, cobalt in pink and sulphur in yellow. Ala261, labelled in red, is the only difference between BtuG2 and BtuG3 binding sites (B), which does not make any interaction.

**Supplementary figure 4.**
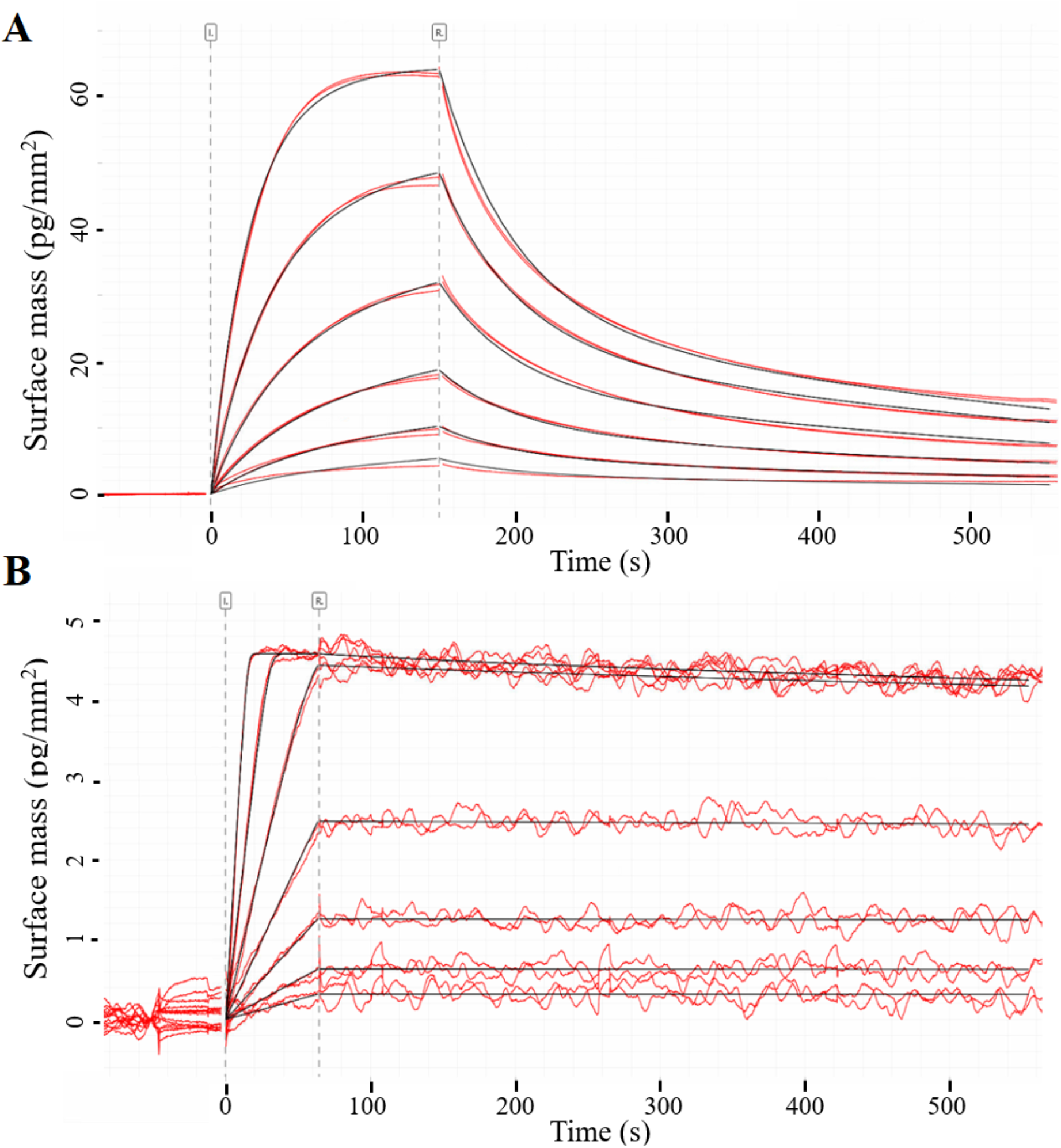
Binding of Cbi and CNCbl to BtuG homologs on a WAVE system PCH-NTA chip. A substrate solution (see Methods) with (A) 6 different concentrations of Cbi at 1.5, 3, 6, 12, 24 and 48 µM (lower to higher surface mass) and (B) 7 different concentrations of CNCbl at 0.08, 0.16, 0.32, 0.64, 1.28, 2.56 and 5.12 nM (lower to higher surface mass) is loaded on a microfluidics chip, pre-loaded with (A) BtuG3 and (B) BtuG2. All concentrations are loaded in technical duplicates. Black lines represent the calculated kinetic models for the experiment, (A) Kd = 38.5 and 3.57 µM with a heterologous ligand model and (B) Kd = 1.03 pM with mass transport limitation of Kt = 3.47 · 10^7^. Double referencing is applied in all experiments against an empty channel and a blank load (see Methods). Dashed lines indicated the start of the sample injection (I) and rinse with running buffer (R).

